# The *MITF* paralog *tfec* is required in neural crest development for fate specification of the iridophore lineage from a multipotent pigment cell progenitor

**DOI:** 10.1101/862011

**Authors:** K. Petratou, S. A. Spencer, R. N. Kelsh, J. A. Lister

## Abstract

Understanding how fate specification of distinct cell-types from multipotent progenitors occurs is a fundamental question in embryology. Neural crest stem cells (NCSCs) generate extraordinarily diverse derivatives, including multiple neural, skeletogenic and pigment cell fates. Key transcription factors and extracellular signals specifying NCSC lineages remain to be identified, and we have only a little idea of how and when they function together to control fate. Zebrafish have three neural crest-derived pigment cell types, black melanocytes, light-reflecting iridophores and yellow xanthophores, which offer a powerful model for studying the molecular and cellular mechanisms of fate segregation. Mitfa has been identified as the master regulator of melanocyte fate. Here, we show that an Mitf-related transcription factor, Tfec, functions as master regulator of the iridophore fate. Surprisingly, our phenotypic analysis of tfec mutants demonstrates that Tfec also functions in the initial specification of all three pigment cell-types, although the melanocyte and xanthophore lineages recover later. We show that Mitfa represses tfec expression, revealing a likely mechanism contributing to the decision between melanocyte and iridophore fate. Our data is consistent with the long-standing proposal of a tripotent progenitor restricted to pigment cell fates. Moreover, we investigate activation, maintenance and function of tfec in multipotent NCSCs, demonstrating for the first time its role in the gene regulatory network forming and maintaining early neural crest cells. In summary, we build on our previous work to characterise the gene regulatory network governing iridophore development, establishing Tfec as the master regulator driving iridophore specification from multipotent progenitors, while shedding light on possible cellular mechanisms of progressive fate restriction.

## Introduction

Pigmentation is a conspicuous feature of animal diversity and has broad importance for behaviour and evolution (reviewed in [1]). Much is known about the development and cell biology of melanocytes but far less is understood about the genetic mechanisms underlying the diversity of pigment cell types in non-mammalian vertebrates. Zebrafish embryos display three neural crest (NC)-derived pigment cells: melanophores, melanin-producing cells homologous to the melanocytes of mammals, and often referred to simply as melanocytes; xanthophores, yellow-orange cells bearing pteridine and carotenoid pigments; and iridophores, shiny cells containing platelets composed of guanine (reviewed in [2], [3]). Defining the fate specification mechanisms of these other cell-types is important for understanding their genetic control and their evolutionary origins.

The progressive fate restriction model proposes that neural crest cells (NCCs) become partially fate restricted as development progresses, giving rise to intermediate partially-restricted progenitors, each of which can generate a number, but not all of the NC derivatives [4]–[8]. Such a model has been strongly supported for the neural derivatives by a single cell transcriptional profiling study in mice, which surprisingly was unable to resolve pigment cell development [9]. Characterisation of the phenotypes of mutants affecting multiple NC derivatives has been widely used to infer the identities and potencies of these progenitors. In the pigment cell field, a progressive fate restriction model has been developed, with both a multipotent chromatoblast and a bipotent melano-iridoblast as identified partially-restricted intermediates ([10] [11]–[14]). Studies of fate-specification mutants further permits elucidation of the gene regulatory networks (GRNs) governing diversification of these precursors ([14], [15]).

To date, examination of zebrafish mutants provides evidence of complex genetic control of pigment cell development from multipotent NCCs [16]. Of the genes affected in these mutants, many have been shown to encode transcription factors that regulate fate specification of pigment cells from NCCs. Such factors may be required by either all three pigment cell lineages (e.g. *colourless/sox10*; [17], [18]) or only a single pigment cell lineage (e.g. *nacre/microphthalmia-associated transcription factor a*/*mitfa*, [19]). Consequently, genetic loss of several transcription factors affects one or more pigment cell types, indicating the existence of shared progenitors. For example, loss of *sox10* function results in lack of all three zebrafish lineages, but also hinders the development of peripheral nervous system components ([20], [21]). Intriguingly, a gene mutation affecting *only* the three pigment cell lineages has not been identified, thus the existence of a tripotent progenitor, exclusively generating chromatophore lineages [10], and the process of pigment cell diversification remain unclear.

As a first step towards understanding the complex GRN governing NCC fate restriction towards pigment cell lineages, it is important to define the key components involved in fate specification of individual lineages. Of the three zebrafish pigment cells, melanocytes are the currently best studied. In this lineage, Sox10, in conjunction with Wnt signalling, is required to activate and maintain *mitfa* transcription ([15], [18], [22]–[24]). Like its mammalian counterpart, MITF, Mitfa has been proven necessary and sufficient to upregulate numerous melanocyte differentiation genes, including those controlling melanin synthesis (e.g. *dct*, *silva* and *tyrosinase*). Mitfa is thus dubbed the ‘master regulator’ of melanocyte fate choice ([15], [19], [23]). However, in this role, Mitfa is supported by Tfap2, especially Tfap2a which acts as a key co-factor [25]. In contrast, Foxd3 mutants show a complex mutant phenotype that includes delayed melanocyte and xanthophore specification and differentiation, ultimately resulting in wild-type cell numbers, and reduced iridophore numbers ([12], [26]–[28]). These phenotypes seem to reflect roles for FoxD3 in both lineage priming [29] and, in certain contexts, repressing melanogenesis promoting the specification of other fates ([12], [13]).

Recent studies of key zebrafish iridophore mutants have begun to define the basic genetics of this cell type. In terms of their differentiation, heightened purine synthesis is central to the development of the guanine crystals that form the reflecting platelets. Thus, *purine nucleoside phosphorylase 4a (pnp4a),* which encodes an enzyme important in the biosynthesis of guanine, is a robust marker of mature iridophores ([13], [14]). Additionally, mutations of several enzymes specific to differentiated iridophores have been shown to disrupt purine biosynthesis [30], while disruption of other proteins was found to impair iridophore survival ([31]–[33]). Furthermore, a signalling pathway crucial to fate specification, proliferation and differentiation of iridophores have been highlighted by mutations in the gene encoding the Leukocyte Tyrosine Kinase (Ltk; [11], [34]) receptor tyrosine kinase, with corroboration from targeted loss of function of its ligand ([35], [36]). In *shady/ltk* mutants fate specification and proliferation of iridophores, but not other pigment cells, are impeded ([11], [34]). Nevertheless, Ltk signalling alone is not sufficient for iridophore specification, since iridophores are eliminated in *sox10* mutants, even though *ltk* expression is strongly detectable by WISH ([11], [14]). The function of Ltk signalling in specification of the iridophore lineage is, thus, likely analogous to that of Wnt signalling in generating melanocytes. This then leaves open the question of whether there is a ‘master’ transcriptional regulator of the iridophore lineage, analogous to the role of Mitfa in melanocytes.

The zebrafish “MiT” (Mitf/Tfe) family consists of six genes [37]; of these, *mitfa* and *tfec* are the only ones expressed in the NC. The distinctive expression pattern of *tfec* in cells along the dorsal and ventral midline in the trunk and tail and two patches over the yolk, is strongly reminiscent of differentiating iridophores [37]. Furthermore, Higdon et al. performed transcriptomic analysis to compare FACS-purified iridophores, melanocytes, and retinal pigment epithelium (RPE), and found *tfec* to be one of several transcription factor genes with enriched expression specifically in the iridophore lineage [38]. Together, these data lead to the hypothesis that Tfec might be the iridophore master regulator, equivalent to Mitfa in developing melanocytes.

In a recent detailed study, we demonstrated that, indeed, *tfec* serves as a robust marker of the iridophore lineage, allowing us to define the major stages of iridophore development [14]. Based on whole-mount *in situ* hybridisation (WISH) studies of *tfec* expression patterns throughout a developmental time-course, and co-expression analysis with both *ltk* and *mitfa*, we concluded that *tfec* is first expressed extensively throughout the premigratory NC progenitors of the trunk and tail at 18 hpf, and then exclusively in the developing iridophore lineage. Focussing on cells in the posterior trunk, we distinguished a subset of *tfec*-positive cells within the premigratory domain that have downregulated the early NCC marker, *foxd3*, but do not express definitive pigment cell markers; we refer to these as early chromatoblasts (*Cbl early*). At approximately 22 hpf, *ltk* and *mitfa* are upregulated in the *tfec*+; *foxd3*-premigratory cells of the trunk ([11], [14], [19]), indicating that they correspond to partially restricted progenitors, capable of generating pigment cells (*Cbl late*). By 24 hpf, *tfec* labels migrating progenitors which co-express *mitfa* and *ltk* markers, and which we consider fate-specified iridoblasts, but which likely retain at least melanocyte potential too (*Ib(sp);* [14]). From 30 hpf, *tfec+* cells co-express *ltk*, but not *mitfa*, and we now consider these to be either definitive iridoblasts (*Ib(df)*, along the dorsal trunk and on the lateral migration pathway at 30 hpf), or mature iridophores (*Iph;* in iridophore locations from 42 hpf onwards) according to their position and state of visible differentiation. Thus, the established iridophore-associated expression pattern of *tfec* during zebrafish development reinforces the hypothesis that Tfec might act as the missing iridophore master regulator, but does not eliminate the possibility of additional earlier functions, either in multipotent early NCCs (eNCCs), or in partially restricted precursors with wider potencies.

In our previous study we used a preliminary assessment of *tfec* mutants to inform our derivation of a core iridophore GRN. Here, we describe in detail the generation and comprehensive characterization of the effects on NC development of mutations in *tfec*. Following examination of the development of a wide variety of NC derivatives in *tfec* mutants, using early and late molecular markers, we conclude that, although neuronal and skeletal derivatives develop normally, specification of *all* pigment cell fates is delayed in homozygous mutants, suggesting a common early requirement for *tfec* in the GRN governing specification of all three chromatophore lineages, and providing support for a common chromatoblast precursor. Finally, our previous work identified the GRN governing maintenance of *tfec* in the iridophore lineage [14]. In the present study, we extend this work to define *tfec* as the iridophore master regulator and, importantly, to identify its upstream regulators in the multipotent premigratory NC, thus placing the transcription factor in context in the NCC specification GRN [39]. Together, these data shed light on the possible mechanism of progressive fate segregation of NCCs, and begin to elucidate the complex role for Tfec, being indispensable for iridophore development, but also playing subsidiary roles in specification of the other two chromatophores derived from the zebrafish NC.

## Results

### *tfec* is a candidate for the iridophore master regulator

As we showed previously [14], *tfec* is co-expressed with the established iridophore marker, *ltk* [11], during iridoblast fate choice and iridophore differentiation. Although Higdon et al showed that *tfec* expression was prominent in the RNA-seq profiles of iridophores, they also detected low levels of expression in purified melanocytes [38]. Here we used WISH to detect *tfec* in individual embryos, following photographic documentation of their individual iridophore patterns, to show definitively its presence in differentiated iridophores (Fig. 1). *tfec* is maintained in differentiated cells (Fig. 1A-D). At these stages, consistent with our previous observations showing no overlap of *tfec* and the melanocyte marker *mitfa* in differentiated melanocytes [14], we do not detect expression in neighbouring differentiated melanocytes occupying the dorsal and ventral stripes (Fig. 1A-D). Likewise, xanthophores, which are widespread under the epidermis of the flanks of the embryos, also do not show detectable *tfec* expression in these WISH studies (Fig. 1A-D). To confirm this, we used the xanthophore lineage marker, Pax7, detected via an immunofluorescence assay combined with simultaneous labelling of *tfec* transcript via WISH (Fig. 1E-G). In conclusion, at the detection threshold of WISH, *tfec* is a consistent marker of mature iridophores, but not of melanocytes nor xanthophores.

**Figure 1.**
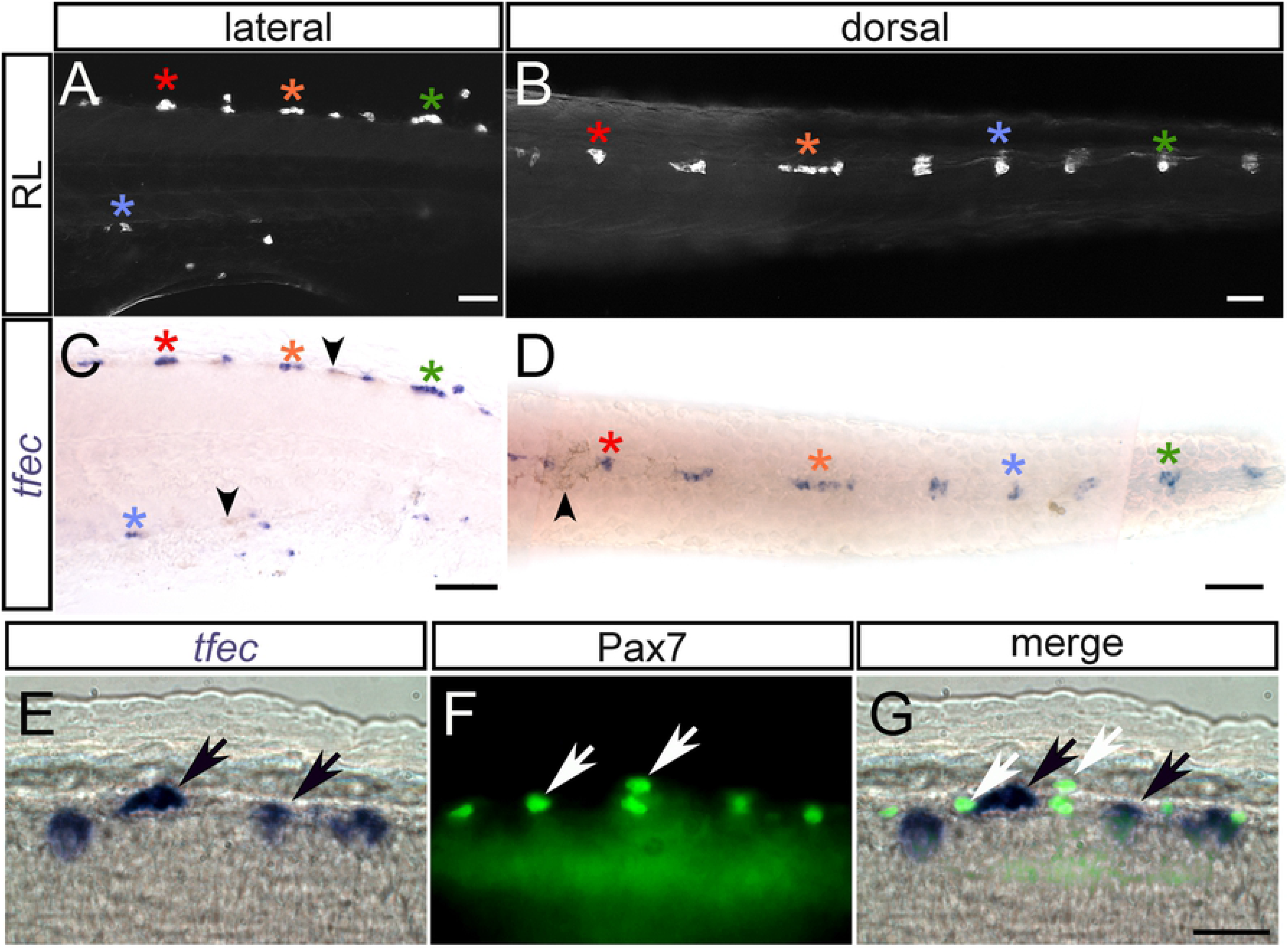
*tfec* is a marker of differentiated iridophores, but not xanthophores. (A-D) Chromogenic WISH reveals *tfec* expression in iridophores at 3 dpf. Iridophores (asterisks) of the posterior trunk/anterior tail are imaged live with reflected light in lateral (A) and dorsal (B) views. (C,D) WISH on the same embryo reveals a pattern of *tfec* expression matching that of the differentiated iridophores (asterisks). Note the absence of expression in the location of the associated melanocytes (residual melanin indicated by arrowheads) (E-G) Expression of *tfec* (E, chromogenic WISH) and of Pax7 (F, immunofluorescence) in differentiated iridophores and xanthophores, respectively, at 48 hpf; note the absence of co-expression of these markers. A,C,E: lateral views; B,D: dorsal views. Head towards the left. Scale bars: A-D, 50 μm; E-G, 20 μm.

### Loss of *tfec* function affects the development of all embryonic pigment cells

To assess the role of *tfec* in development, we induced mutations in *tfec* using CRISPR/Cas9 (Fig. 2), selecting a target in the seventh exon of the gene, which encodes the second helix of the transcription factor’s helix-loop-helix dimerization domain. Our two laboratories independently generated identical 6 base pair deletions (the *tfec^ba6^* and *tfec^vc58^* alleles) in two different wild-type strains, WIK and NHGRI-1, in addition to frameshifted alleles (Fig. 2A). We reasoned that in this region of the gene it was likely that even indels that retained the correct reading frame (i.e., multiples of three) would likely be deleterious, because they would alter spacing of key residues and surfaces within this helix. Indeed, all of the generated alleles, when made homozygous, resulted in elimination of differentiated iridophores from the dorsal, ventral and yolk sac stripes, as well as from the lateral patches of the embryo (Fig. 2F-I). In addition, iridophores were absent from the dorsal head (Fig. 2G,I) and the eye (Fig. 2F-I) of homozygous mutants. Moreover, both injected (G0) fish raised to adulthood, as well as a single ‘escaper’ surviving F1 adult carrying biallelic frameshift mutations, lack iridophores in patches or in whole (Fig. S1 A-C). All results presented here were produced using either the *tfec^ba6^* or the *tfec^vc60^* alleles, unless stated otherwise.

**Figure 2.**
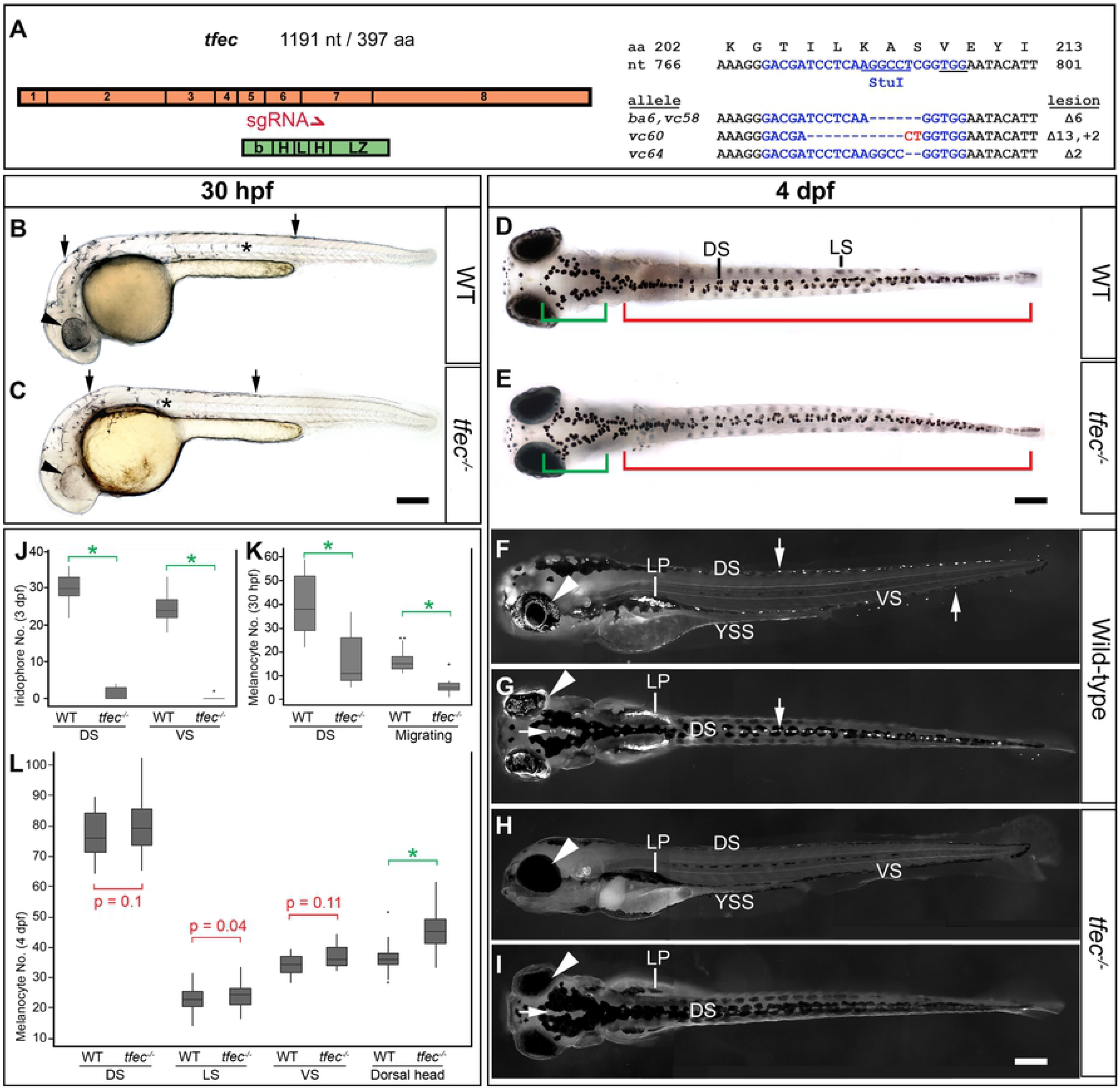
Embryonic pigmentation is affected in *tfec* mutants. (A) Schematic showing the distribution of the 8 exons of *tfec* (orange) in relation to the functional basic helix-loop-helix-leucine zipper domains (green) of the transcription factor. The red arrow indicates the position along both the gene and protein sequences targeted for mutagenesis by the CRISPR/Cas9 system. Included are the targeted WT *tfec* DNA/amino acid sequence (blue, with PAM underlined), and the sequences of the examined mutant alleles, with the corresponding molecular lesions (dashes for deleted nucleotides, red font for insertions). At 30 hpf, compared to WT siblings (B), *tfec* mutants (C) have reduced melanisation of the RPE (arrowheads), and reduced melanocytes both along the dorsal trunk (arrows) and on the migratory pathways (asterisks). (D-I) Pigment cell phenotypes at 4 dpf. In embryos treated with melanin-concentrating hormone (MCH) to facilitate their quantitation, the number of trunk and tail melanocytes is not significantly altered along the dorsal, ventral or lateral stripes (D,E, red region; quantitated in L), however there is a statistically significant increase in the number of melanocytes located on the dorsal head (D,E, green region; quantitated in L). Imaging live embryos under reflected light reveals a striking lack of iridophores in *tfec* mutants (H,I) compared to their WT siblings (F,G) along the dorsal (downward arrow), ventral (upward arrow), and yolk sac stripes, as well as overlying the eye (arrowhead). Iridophores are also absent on the dorsal head (G,I, horizontal arrow) and the lateral patches (LP). (J-L) Quantitation of pigment cell phenotypes in *tfec* mutants. (J) Quantitation of differentiated iridophores at 3 dpf confirms a prominent lack of iridophores along the dorsal and ventral stripe of *tfec* mutants. (K) Quantitation of melanocytes along the dorsal trunk and migratory pathways at 30 hpf reveals a 60% reduction in both regions in *tfec* mutants with respect to WT siblings. sgRNA, small guide RNA; DS, dorsal stripe; VS, ventral stripe; LP, lateral patches; YSS, yolk sac stripe. (B-C,F,H): lateral views. (D-E,G,I): dorsal views. Head towards the left. Scale bars: 200 μm. (J-L): spots signify outlier values; *, p-value < 10^−9^ using t-test.

Quantification of iridophore numbers along the dorsal and ventral stripes of live embryos at 3 dpf illustrates the severity of the phenotype, with only very rare escaper iridophores present in homozygous *tfec^ba6^* mutant embryos (Fig. 2J; Table S1). The numbers of differentiated cells in heterozygous mutants are not significantly different from those in wild-type (WT) siblings (Table S1). The iridophore phenotype could be successfully rescued via injection of a Tol2 transposon-based plasmid containing 2.4 kb of the *tfec* promoter [40], guiding tissue-specific expression of full-length *tfec* (Fig. 3). Although the WT number of iridophores was not recovered, almost half of the injected mutant embryos presented with rescue of iridophores either on the eye, the trunk, or both of those domains (Fig. 3E). Successful iridophore rescue was also visible in injected fish raised to adulthood (Fig. S1 D).

**Figure 3.**
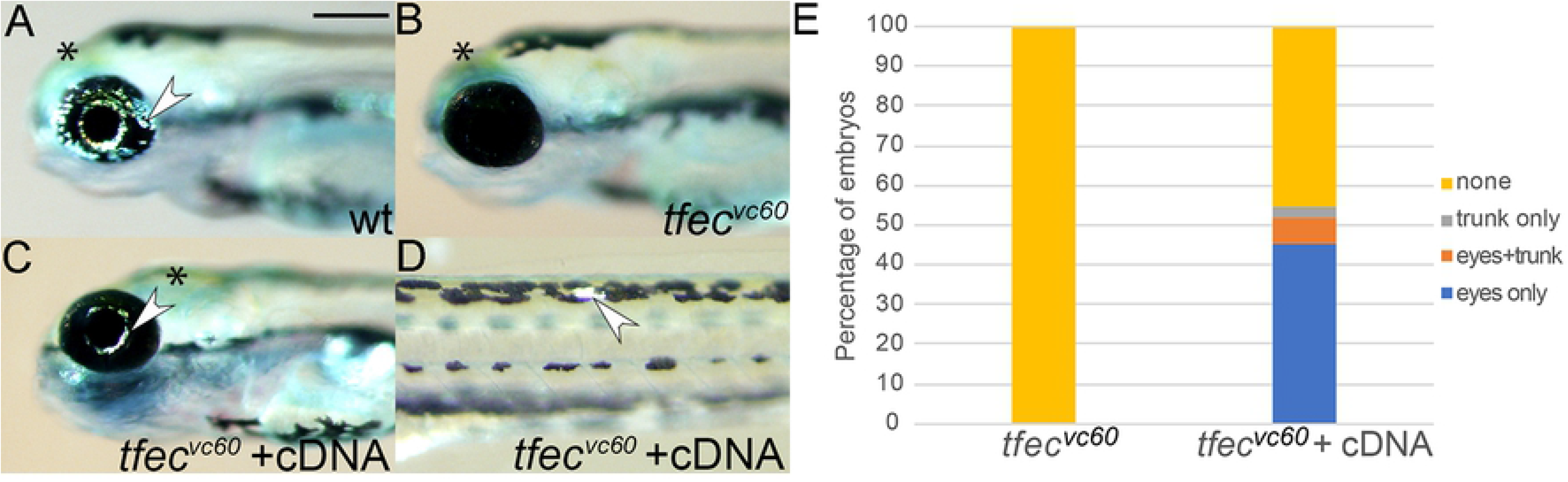
Injection of *tfec* cDNA can rescue the mutant phenotype. Differentiated iridophores (arrowheads) are abundant on the eye of WT embryos (A), but completely absent from the eye of a *tfec^vc60^* mutant sibling (B) at 4 dpf. Co-injection with Tol2 transposase of a construct where the *tfec* promoter drives transcription of the *tfec* cDNA sequence, leads to rescue of iridophores (arrowheads) on the eye (C) and trunk (D) of *tfec^vc60^* mutants. (E) Quantitation of rescue efficiency. Approximately 45% of mutants displayed eye rescue, 3% showed rescue in the trunk only and 6% showed rescue both in the eyes and trunk (n = 62). By contrast, iridophores were never observed in uninjected *tfec^vc60^* sibling larvae (n = 58). Lateral views, heads towards the left. Scale bar: 200 μm.

In previous work, we showed that *tfec* is present in multipotent premigratory NCCs, which do not yet express definitive pigment cell markers (early NCCs, early Cbls; [14]). We further demonstrated that during early stages of specification and migration of NC derivatives, *tfec* expression transiently overlaps with that of *mitfa* in specified, but not definitive, iridoblasts (Ib(sp)). Here we report that melanogenesis is delayed in 30 hpf homozygous *tfec* mutant embryos when compared to WT or heterozygous siblings, supporting a functional role for Tfec during its transient expression in melanoblasts. Specifically, we observed a significant reduction in the numbers of differentiating melanocytes along the dorsal trunk, and the medial and lateral migration pathways (Fig. 2B,C,K; Table S1). Melanocyte development recovers, and by 4 dpf mutant embryos present with the same number of melanised cells along their trunk as their WT or heterozygous siblings (Fig. 2D,E,L; Table S1). Furthermore, we see strikingly reduced melanisation of the retinal pigmented epithelium (RPE) of homozygous mutant embryos at 30 hpf, compared to WT or heterozygous siblings (Fig. 2B,C), suggesting an analogous role in these brain (not NC)-derived melanocytes. Surprisingly, we observed a mild, yet consistent and statistically significant increase in differentiated melanocytes on the dorsal head of mutant embryos at 4 dpf (Fig. 2D,E,L; Table S1).

The delayed melanogenesis phenotype in these mutants might result from delayed specification of melanoblasts, or from slowed differentiation of normally specified melanoblasts. To distinguish between these two possibilities, we performed chromogenic WISH at 24, 30 and 48 hpf to detect expression of the melanocyte master regulator, *mitfa*. Strikingly, *mitfa* expression was restricted to premigratory late Cbls in *tfec* mutants, whereas *mitfa*-positive melanoblasts occupied the medial and lateral migratory pathways in WT and heterozygous siblings (Fig. 4M,N). At 30 hpf, the delay was still detectable. The trunk was occupied by a relatively small number of *mitfa*-positive NC derivatives, whereas in the tail of mutants cells had still not entered the migratory pathways (Fig. 4O,P). Consistent with the live phenotype, *mitfa* expression in mature melanocytes at 48 hpf was unaffected in the trunk and tail of *tfec* mutants, compared to WT siblings (Fig. 4Q,R). This early retardation of *mitfa* expression, coupled to absence of *ltk* expression (Fig. 4S,T; [14]), but normal migration of neural derivatives along the medial pathway (Fig. 5G,H), suggested that specification of the *mitfa*+; *tfec*+ Ib(sp) [14] from late Cbl, was hindered in the absence of functional Tfec. In addition, these data support our previous suggestion [14] that these Ibl(sp) retain melanocyte potential (i.e. they can be considered both specified melanoblasts as well as specified iridoblasts), and show that melanocyte fate specification is delayed in the *tfec* mutant. The subsequent recovery of normal melanocyte numbers makes clear that compensatory factors allow melanoblasts to be specified, albeit with a short delay.

**Figure 4.**
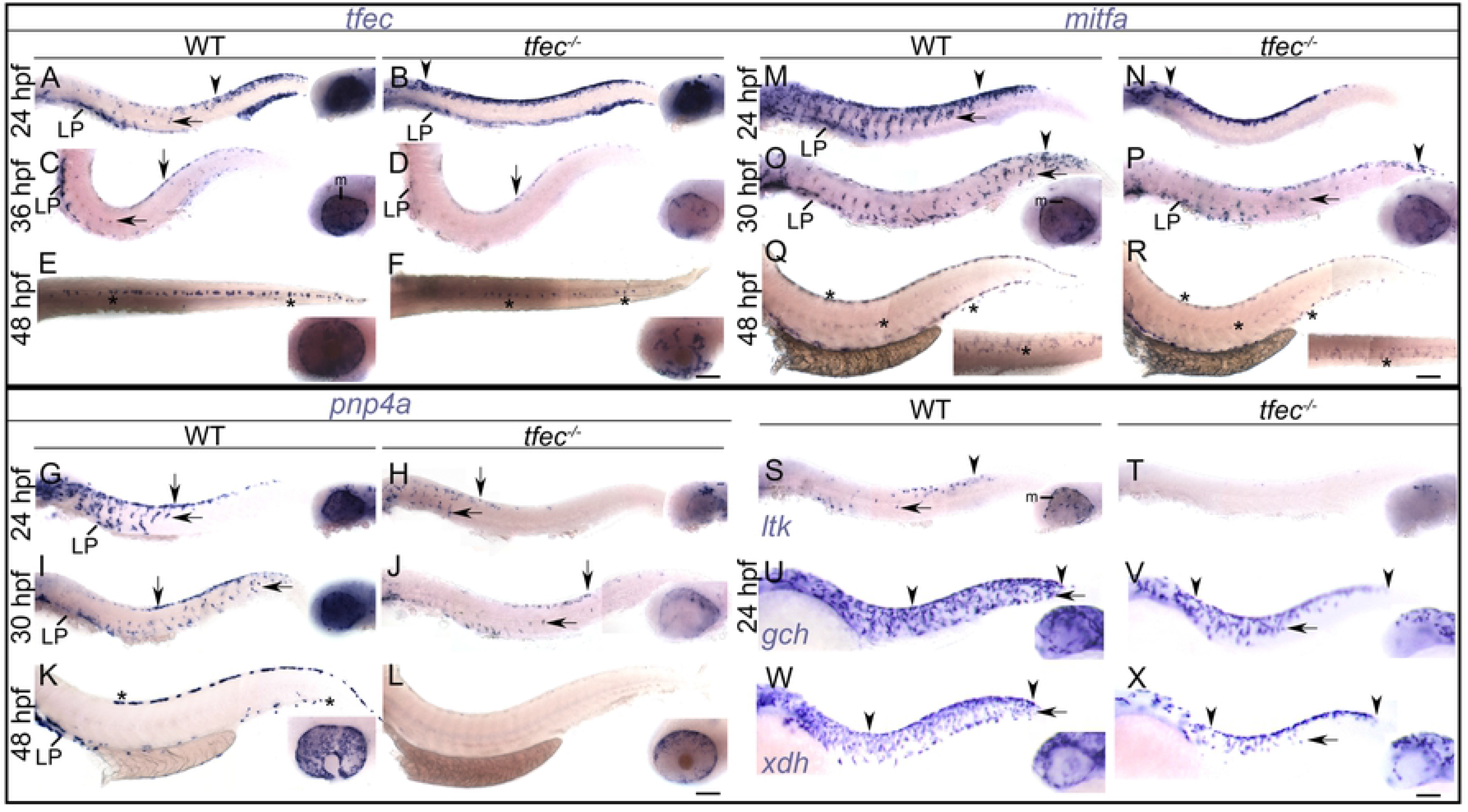
Specification of melanocytes, iridophores and xanthophores is delayed in *tfec* mutants. (A-F) Generation of *tfec*-positive Ib(sp) from late Cbls requires Tfec. At 24 hpf, *tfec* mutants present with anterior expansion of the *tfec*+ premigratory NC domain (A,B, arrowheads) and lack of medially migrating, Ib(sp) progenitors (arrows). At 36 hpf, the number of dorsal and migrating *tfec*+ Ib(df) (arrows), as well as those located in the lateral patches is reduced in mutants, compared to WT or heterozygous siblings (C,D). At 48 hpf, a reduced number of cells (asterisks) express *tfec* along the dorsal stripe of mutant embryos, compared to WT or heterozygous siblings (E,F). *tfec* mutant embryos were distinguishable after WISH by the lack of RPE melanisation (C-F, insets). *tfec*+ iridophore lineage cells overlying the RPE are noticeably reduced in *tfec* mutants compared to WT siblings (A-F, insets). (G-L) *pnp4a* expression in Ib(sp) requires Tfec. At 24 hpf, the number of dorsally located, and of partially fate restricted, migrating chromatoblasts (arrows) expressing *pnp4a* is reduced in *tfec* mutants (G,H). Expression overlying the eye is also affected (G,H, insets). This reduction is still prominent at 30 hpf (I,J), when staining is also visibly affected in the lateral patches. *pnp4a* expression is undetectable in differentiated iridophore positions (K, asterisks) in *tfec* mutants at 48 hpf (L). *pnp4a*+ iridophore lineage cells overlying the RPE are noticeably reduced in *tfec* mutants compared to WT siblings (G-L, insets). (M-R) Mb(sp) generation from the late Cbl is delayed in *tfec* mutants. At 24 hpf, *mitfa* marker expression is restricted to an increased number of premigratory Cbls (arrowheads) and is undetectable in migrating Mb(sp) (arrows) in *tfec* mutants (M,N). At 30 hpf, the numbers and distributions of *mitfa*-positive late Cbls (arrowheads) are indistinguishable between *tfec* mutants and WT or heterozygous siblings (O,P), but the delay in Mb(sp) migration remains distinguishable towards the tail (arrows). *tfec* mutant embryos lack RPE melanisation (O,P, insets). At 48 hpf, *mitfa* expression in mature melanocytes (asterisks) is indistinguishable between *tfec* mutants and WT or heterozygous siblings (Q,R). (S, T) *tfec* mutants lack *ltk* expression in premigratory late Cbls (arrowhead), in migrating Ib(sp) (arrow) and in iridoblasts of the eye (insets). (U-X) Xanthoblast (Xbl(sp)) specification from Cbls is delayed, as indicated by WISH against two lineage markers, *gch2* (U,V) and *aox5* (W,X). At 24 hpf, laterally migrating Xbl(sp) (arrows) are restricted to more anterior regions and precursors located in the head are reduced in *tfec* mutants (V,X) compared to their WT siblings (U,W). All *tfec* mutant panels show the *tfec^ba6^* allele except (V,X) which show the *tfec^vc60^* allele. LP, lateral patches; m, melanisation. (A-D,G-X): lateral views. (E-F, insets of Q-R): dorsal views. Head towards the left. Scale bars: 100 μm.

**Figure 5.**
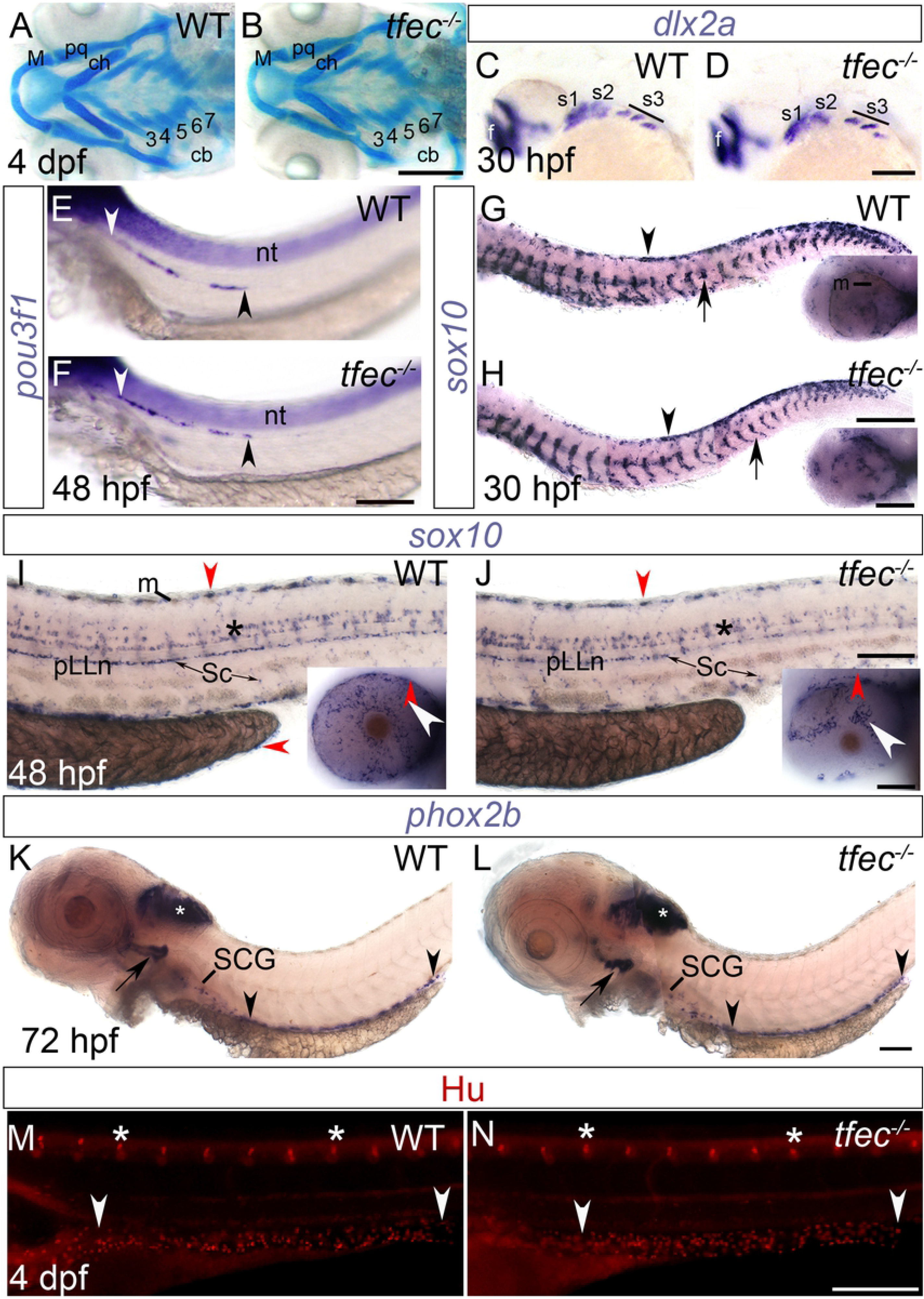
Development of skeletal and neural NC derivatives is unaffected in *tfec* mutants. (A,B) The cranial cartilage at 4 dpf is unaffected in *tfec* mutants, versus WT siblings, as indicated by Alcian blue staining. Numbered 3-7 are the positions of the branchial arches. (C,D) At 30 hpf, *dlx2a* expression, detected by WISH, shows that formation of the three streams (s1-s3) of migrating cranial NCCs is unaffected in *tfec* mutants. Staining is also indistinguishable in the forebrain (f). (E-N) Peripheral nervous system derivatives develop normally in *tfec* mutants. (E,F) *pou3f1* expression analyses at 48 hpf show that glial progenitors on the posterior lateral line (pLLn; area between arrowheads) develop normally in *tfec* mutants. (G,H) *sox10* staining at 30 hpf indicates no observable alterations in the migration of specified neural progenitors through the medial pathway (arrows) in *tfec* mutants, compared to WT siblings. Likewise, the number and distribution of *sox10-*positive premigratory NC progenitors (arrowheads) is unaffected. *tfec* mutants were identified by lack of RPE melanisation (G,H, insets). (I,J) At 48 hpf, *sox10* expression analysis showed indistinguishable numbers and distribution of Schwann cells (Sc) occupying the pLLn and spinal nerves. Likewise, oligodendrocyte progenitors throughout the CNS appear normal in their specification, numbers and migration (asterisks). *sox10* expression is detectable in iridophore positions (red arrowheads) and in eye iridophores (insets, white arrows), that are strongly affected in homozygous *tfec* mutants. (K,L) *phox2b* expression, detected by WISH, at 72 hpf. The formation and the extent of migration of enteric nervous system progenitors (region between arrowheads) are indistinguishable between *tfec* mutants and WT siblings. Likewise, expression in the earliest differentiating region of the sympathetic ganglia chain, the superior cervical ganglion (SCG), is unaffected. *phox2b* expression is also indistinguishable in the hindbrain (white asterisks) and in placode-derived neuronal progenitors in the cranial ganglia associated with the branchial arches (arrows). (M,N) At 4 dpf, the DRG (asterisks) and enteric neurons (arrowheads) number and positioning, as revealed by immunofluorescent detection of Elav1/Hu is indistinguishable between *tfec* mutant embryos (N) and WT siblings (M). (B,D,F,N): *tfec^vc60^* allele, (H,J,L): *tfec^ba6^* allele. cb, ceratobranchials; ch, ceratohyal; M, Meckel’s cartilage; nt, neural tube; pq, palatoquadrate. (A-B): dorsal views. (C-N): lateral views. Head towards the left. Scale bars: (A-F, M-N): 200 μm; (G-L): 100 μm.

We set out to investigate the early specification events that lead to the observed pigmentation phenotypes using WISH studies of iridophore markers. In homozygous *tfec* mutants, expression of the differentiated iridophore marker, *pnp4a* ([13], [14]), was undetectable at 48 hpf (Fig. 4K,L), consistent with complete lack of differentiated iridophores in *tfec* mutants. At both 24 hpf and 30 hpf, *pnp4a* was expressed in a notably reduced number of cells along the dorsal trunk, the migratory pathways and overlying the eye (Fig. 4G-J). Maintenance of *pnp4a* in a small subset of cells at these earlier stages of chromatoblast specification was attributed to Mitfa-dependent activation ([14]; Fig. 4G-J). Notably, the early reduction of *pnp4a* expressing cells in the eye and trunk of homozygous *tfec* mutants is consistent with a defect in generating the *pnp4a*+ Ib(sp) in these embryos. We assessed expression of *tfec* itself in *tfec* mutant embryos, using our *tfec^ba6^* allele (Fig. 2A). At 24 hpf, presumptive homozygous mutants maintained *tfec* expression along the premigratory NC domain, even in anterior regions, indicating failure of a subset of NC derivatives to become fate specified, since in WT embryos fate specification to non-iridoblast fates is accompanied by loss of *tfec* expression in the majority of derivatives (Fig. 4A,B). Furthermore, *tfec* expression was undetectable in the medial migration pathway, consistent with the absence of *tfec*-positive Ib(sp). At 36 hpf, the number of *tfec*-positive cells identified as Ib(df) [14] was strongly reduced in *tfec* mutant embryos compared to their siblings (Fig. 4C,D), consistent with *tfec* function being fundamental for iridophore specification. Intriguingly, complete lack of differentiated iridophores (Fig. 2F-I) was not accompanied by corresponding total elimination of *tfec* expression at 48 hpf (Fig. 4E,F; [14]). As the remaining *tfec*-positive cells do not express other iridophore markers, such as *pnp4a* (Fig. 4L) or *ltk* [14], we hypothesise that these correspond to early partially-restricted NC derivatives, perhaps early Cbls. Finally, we examine *ltk* expression in *tfec* mutants. *ltk* expression was completely lacking on the medial migration pathway in homozygous mutants at 24 hpf (Fig. 4S,T; [14]), consistent with an early defect in iridophore specification. However, we note that *ltk* expression is also missing in the premigratory Cbl domain, indicating a much earlier role for Tfec, in specification of the Cbl(late) from Cbl(early).

The *tfec* mutant embryos did not show obvious changes in the number and distribution of mature xanthophores (Fig. 3B). However, examination of early xanthophore specification markers by WISH showed that the developmental delay in producing melanoblasts is also true for generation of xanthoblasts (Fig. 4U-X). Specifically, both *aox5-* and *gch2-*positive cells appeared less abundant along the lateral migration pathway at 24 hpf. Delay in the expression of these two genes was also noted in the head of *tfec* mutant embryos, compared to WT siblings (Fig. 4U-X, insets). In conclusion, generation of melanoblasts, iridoblasts and xanthoblasts from multipotent NCCs is delayed in the absence of functional Tfec, pointing to an unexpectedly wide role in the specification of all chromatophore fates.

### Loss of *tfec* function does not affect the development of non-pigment NC derivatives

Given the unexpected role in non-iridophore pigment cells, we then asked whether loss of *tfec* function affected non-pigment NC derivatives. To examine cartilage development, we stained *tfec* mutant embryos and WT siblings with Alcian Blue but did not note any visible phenotypic distinction in homozygous mutants (Fig. 5A,B). To assess neural fates, we used a series of standard markers. The number and distribution of dorsal root ganglion (DRG) sensory neurons, as labelled by anti-Hu immunofluorescence, was unaffected by loss of *tfec* function (Fig. 5M,N). Both our anti-Hu assays and traditional WISH staining for *phox2b* expression (Fig. 5K,L) indicated that development of the NC-derived enteric neurons and enteric and sympathetic progenitor cells remained unaffected in the absence of functional Tfec. Moreover, the number and patterning both of mature Schwann cells, normally residing on the posterior lateral line nerve along the horizontal myoseptum, and of satellite glial cells associated with the DRGs, remain unaffected in homozygous *tfec* mutants, as shown by WISH staining for *sox10* at 48 hpf (Fig. 5I,J). Likewise, developing oligodendrocytes in the CNS are unaffected in their numbers and distribution.

To examine whether specification of either ectomesenchymal or neural derivatives from multipotent progenitors is delayed in *tfec* mutants, similar to non-iridophore pigment cell derivatives, we conducted additional WISH experiments at 30 hpf, when relevant early specification markers are detectable. We thus showed that expression of both *dlx2a* in migrating cranial NCCs (Fig. 5C,D), and of *pou3f1* (previously *oct6*) in migrating precursors of the pLLn appears (Fig. 5E,F), appeared normal in *tfec* mutants, compared to known WT siblings. Furthermore, detection of *sox10* expression in batches containing WT, heterozygous and *tfec* mutant embryos failed to detect alterations in either the distribution or the abundance of glial progenitors migrating along the medial pathway (Fig. 5G-H). Furthermore, these *in situs* provide evidence that premigratory NCCs appear to be present in the normal numbers and show unaltered timing of loss of *sox10* expression (Fig. 5G,H). Note that in assays aiming to detect *sox10* transcript at 30 hpf, we were able to confirm the presence of homozygous mutant embryos based on reduced melanisation of the RPE (Fig. 5G,H, insets). For experiments from 48 hpf onwards (Fig. 5I-L), homozygous mutants were processed separately from their siblings.

In summary, although *tfec* expression is prominent in the majority of, if not all, early NCCs ([14], [37]), we could not detect any changes at any stage of the development of NC-derived neurons, glial cells and skeletal components. This suggests that although *tfec* transcript is present at these early stages, Tfec is uniquely required for pigment cell fate specification, and essential for iridophore fate specification.

### *tfec* is downstream of NC specifier genes in the NC progenitor GRN

To determine the upstream regulators of *tfec* in premigratory early NCCs and early Cbls of the dorsal trunk, we conducted WISH experiments on single and double mutants for the important vertebrate NC specifier genes *foxd3*, *sox9b*, *sox10* and *tfap2a* [39]. Interestingly, at 18 hpf, all embryos from crosses of *foxd3*, *sox10* and *tfap2a* mutant carriers showed identical expression of *tfec*, strongly suggesting that early expression of *tfec* is not strictly dependent upon any one of these genes (Table S2). In contrast, in 18 hpf presumed homozygous *sox9b* mutants, *tfec* expression does not extend towards the tail as far as in WT or heterozygous siblings (Fig. 6K,L; Table S2), which is likely attributable to delayed specification of early NCCs upon loss of *sox9b* function. At 24 hpf *tfec* expression in WT embryos is gradually downregulated from the majority of premigratory NCCs of the trunk in an anterior to posterior manner, strongly persisting only in Ib(sp) [14]. However, we observed a persistence of *tfec* expression in the anterior premigratory NC domain in *sox10*, *sox9b*, *tfap2a* and *foxd3* mutants at this stage, consistent with retained premigratory late Cbls (Fig. 6A-D,M-N). This persistence differed in severity and duration between different mutants, but homozygous mutants of each genotype show highly consistent phenotypes across experimental replicates. Specifically, as was previously reported for a single time point [14], in *sox10* mutants, where NC derivatives fail to become specified and to enter the migration pathways ([11], [18]), *tfec* expression is maintained in trapped late Cbls, extending to the hindbrain/trunk boundary (Fig. 6A,B). Our results show that, at all time-points, *tfec*-positive premigratory progenitors persist in homozygous mutants (identified by their lack of *tfec* expression in Ib(sp) positions), until 36 hpf (Fig. 6A-B,E-F,I-J). In each of *sox9b*, *tfap2a* and *foxd3* homozygous mutants at 24 hpf, *tfec*+ premigratory NCCs are detectable along the dorsal trunk, but not reaching the hindbrain/trunk boundary as in *sox10* homozygous mutants (Fig. 6C,D,N; Fig. S2).

**Figure 6.**
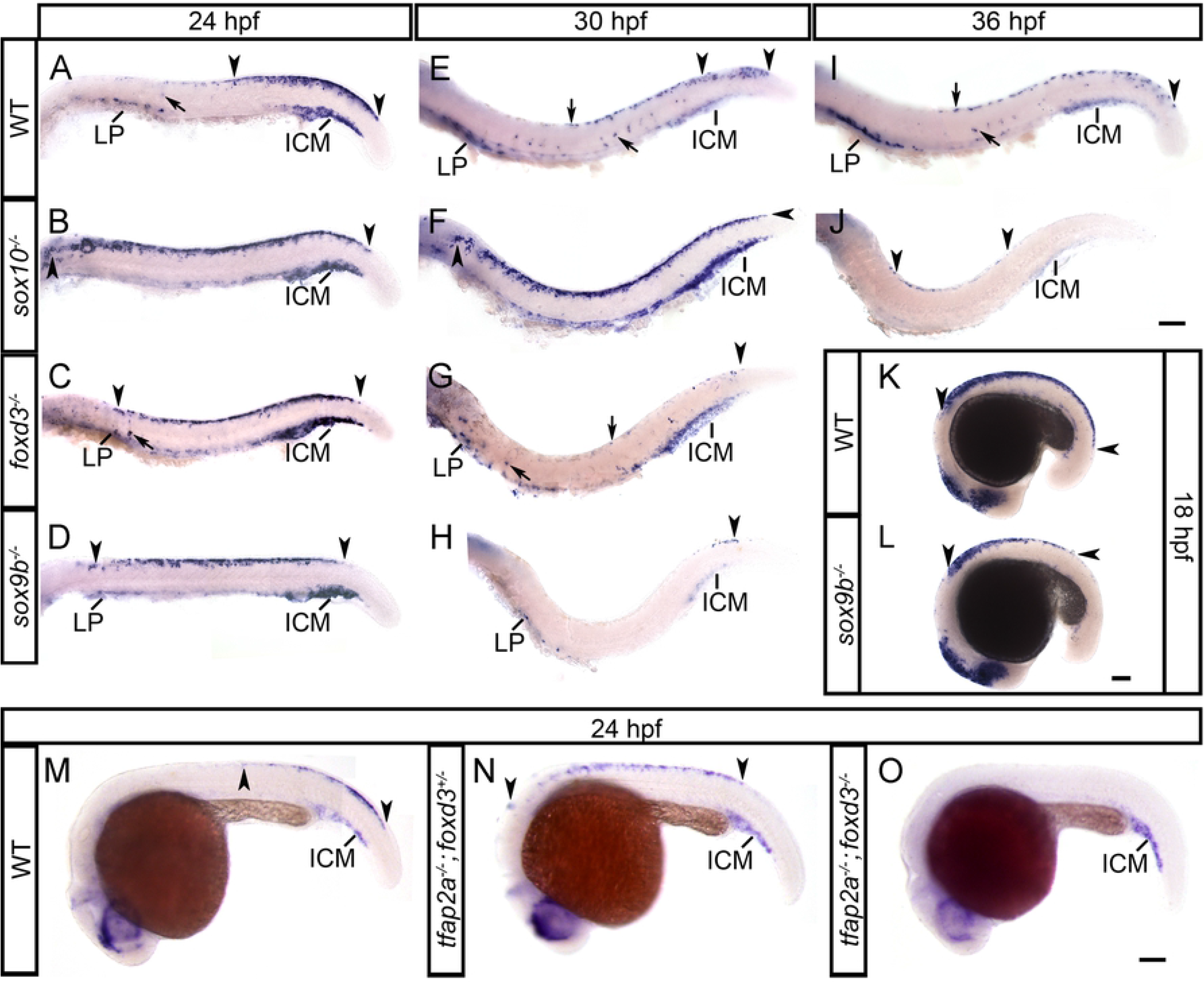
*tfec* is a member of the GRN functioning to specify multipotent NCCs, following their induction. In *sox10*, *foxd3* and *sox9b* mutants (B,C,D) at 24 hpf, *tfec*-positive late Cbls are trapped in the premigratory domain along the trunk, whereas in the WT (A) they are restricted to the dorsal tail (regions between arrowheads). Consistent with failed or delayed development of NC progenitors, migrating (arrow) and LP-located Ib(sp) (A) are reduced (C,D) or completely absent (B). The anterior expansion of the progenitor domain is more pronounced in *sox10* mutants (B). Initiation of expression in eNCCs, detected by posterior-most boundary of expression (posterior arrowhead), is normal in *sox10* (B) and *foxd3* (C) mutants, but is perceptibly delayed in *sox9b* mutants (D); this effect is also prominent at 18 hpf (L). In 30 hpf WT embryos (E) *tfec* transcript is detectable in premigratory late Cbls of the posterior tail (region within arrowheads). This domain still shows a dramatic expansion in *sox10* mutants (F, arrowheads), while *foxd3* (G) and *sox9b* (H) mutants no longer present with trapped progenitors. Instead, very few *tfec*+ cells are detectable in the dorsal tail (arrowheads), and there is a prominent reduction in Ib(df) (arrows, LP). (I) In WT embryos at 36 hpf, *tfec* is still expressed in premigratory Cbls in the posterior-most tail (arrowhead), as well as Ib(df) (arrows). In *sox10* mutants (J) trapped premigratory progenitors (region within arrowheads) are reduced, but still visible. *tfec* expression in the premigratory NC domain (region within arrowheads) is anteriorly shifted in single *tfap2a* mutants (N), compared to WT siblings (M) at 24 hpf, but completely eliminated in double *tfap2a*;*foxd3* mutants (O). *tfec* expression is invariably detectable in the intermediate cell mass from 18 hpf to 36 hpf, in WT and mutant embryos (A-O). ICM, intermediate cell mass; LP, lateral patches. Lateral views, head towards the left. Scale bars: 100 μm.

As members of the same SoxE group, it is not surprising that *sox10* and *sox9b* have been shown to be functionally redundant in DRG sensory neuron development [20]. We asked whether this might also be true for *tfec* expression in early or late Cbls. Examination of embryo batches containing *sox10;sox9b* double mutants at 18 hpf did not reveal elimination of *tfec* expression in any of the assessed embryos (Table S2). However, *tfec* expression was completely eliminated from the NCC progenitor domain of genotyped *tfap2a;foxd3* double mutants (Fig. 6M-O), suggesting that both these genes act together to upregulate *tfec* expression in premigratory NCCs of the trunk. This effect was not observed in genotyped siblings, heterozygous for one or both alleles nor those homozygous for a single mutant allele. In conclusion, our data show key roles for *foxd3* and *tfap2a* in establishing expression of *tfec* in premigratory multipotent NCCs.

### Mitfa represses *tfec* during melanocyte development

Both *tfec* and the melanocyte master regulator, *mitfa*, are transiently expressed in the multipotent NCC domain, in late Cbls, as well as in Ib(sp). We conducted WISH to assess the effects of loss of *mitfa* function on *tfec* expression. Interestingly, presumed homozygous *mitfa* mutants show ectopic expression of *tfec* along the dorsal trunk and in NC derivatives along both the medial and lateral migration pathways (Fig. 7A-D). This pattern of *tfec* expression in *mitfa* mutants resembles WT *mitfa* expression in developing melanoblasts, therefore it is likely that Mitfa represses *tfec* expression in NCCs that become biased towards the melanocyte lineage. This effect persists at 30 hpf and is also observed by a complementary, and more sensitive fluorescent W*ISH* technique, RNAscope; we see persistence of *tfec* expression in cells migrating through the lateral pathway (Fig. 7E-F’). These cells co-express the definitive lineage marker, *ltk* ([11], [14]), suggesting that they correspond to specified iridoblasts.

**Figure 7.**
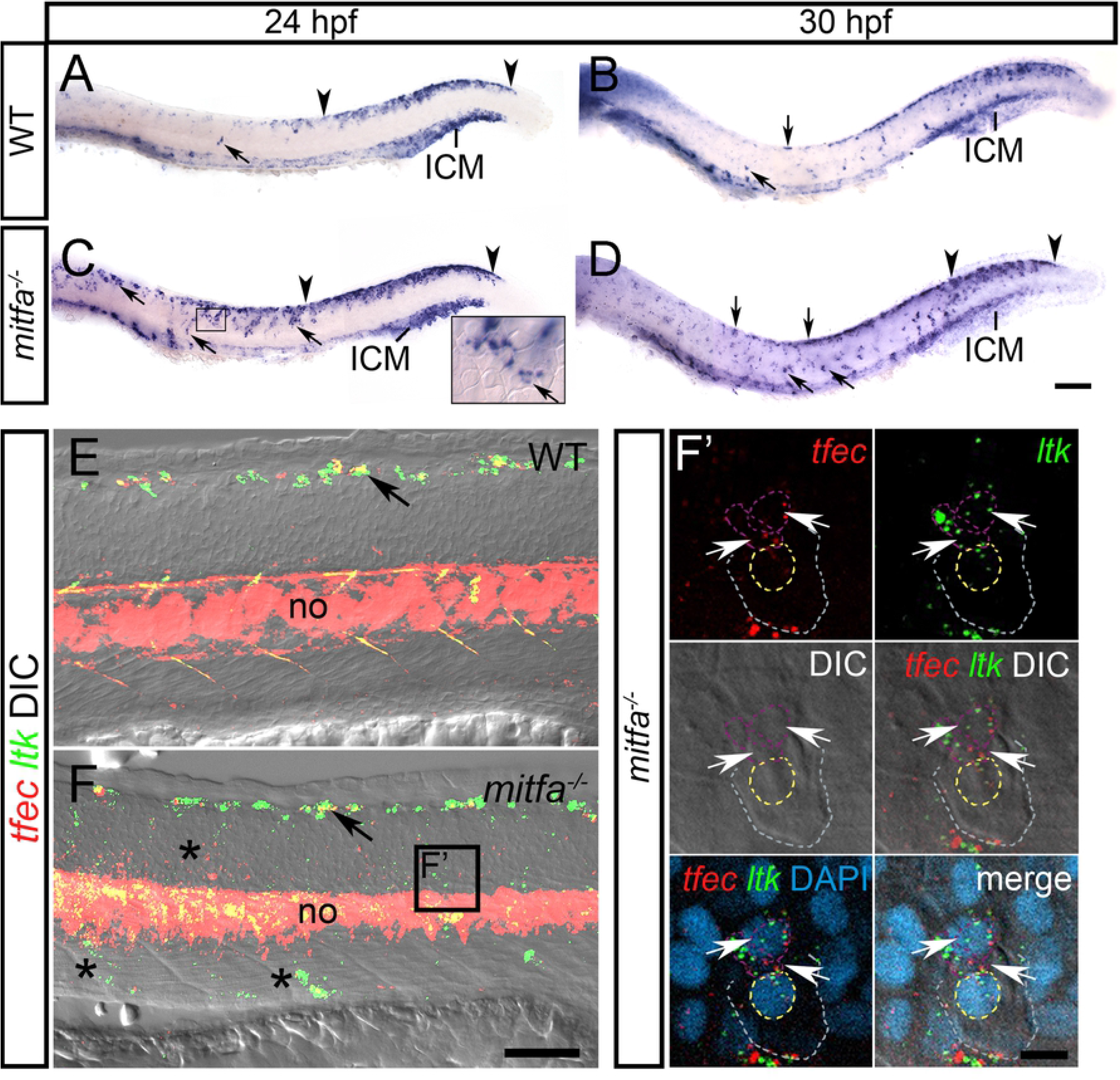
Mitfa represses *tfec* expression in melanoblasts. In WT embryos at 24 hpf (A) and at 30 hpf (B), *tfec* expression is detectable in Ib(sp) and Ib(df) (arrows), respectively, and in the posteriorly regressing early NCC/Cbl domain of the dorsal posterior trunk and tail (region within arrowheads), but expression is undetectable in the lateral migration pathway. At 24 hpf (C) and at 30 hpf (D), *mitfa* mutants present with an increased number of *tfec*-positive cells (arrows) along the dorsal trunk, as well as in the medial and lateral (C, inset) migratory pathways. RNAscope (E,F) performed on *mitfa* mutants at 30 hpf shows an increased number of *tfec*-positive cells compared to the WT on the migration pathways (F, asterisks). Arrows indicate iridoblast precursors along the dorsal trunk. (F’) RNAscope reveals that ectopic *tfec*-positive cells migrating on the lateral migration pathway of *mitfa* mutants (below the epidermis; grey and yellow dashed lines indicate the periphery and nuclear boundary, respectively, of overlying keratinocytes) co-express *ltk* (arrows; nuclei indicated by purple dashed lines), thus likely correspond to Ib(sp). ICM, intermediate cell mass; LP, lateral patches; no, notochord. Lateral views, head towards the left. Scale bars: A-D: 100 μm; E,F: 50 μm; F’: 10 μm.

## Discussion

Our previous work established *tfec* as a marker during NC development and iridophore fate choice ([14], [37]). Interestingly, *tfec* was found to be co-expressed with *mitfa* in Ib(sp) cells, proposed to be able to at least give rise to melanocytes and iridophores, but not in mature melanocytes. Thus, Ib(sp) should also be considered as specified melanoblasts. The first aim of the present study was to determine whether iridophores are indeed the only embryonic differentiated pigment cell expressing *tfec*; to this end, we conducted analyses showing that *tfec* expression is maintained at detectable levels only in mature iridophores, but neither in melanocytes nor in xanthophores.

Considering the strong sequence conservation between Tfec and the melanocyte master regulator, Mitfa, we next asked whether *tfec* might have a function in iridophores analogous to that of *mitfa* in melanocytes; i.e. as the master regulator of the iridophore lineage. We generated mutations in *tfec* using a CRISPR/Cas9 approach, obtaining several alleles which displayed essentially identical phenotypes. Although *tfec* mutants have been generated independently [41], that report did not examine NC-related defects, focusing instead on deficiencies in hematopoiesis. As with the CRISPR-generated exon 3 allele reported by Mahony and colleagues, our mutants fail to inflate the swim bladder (which, along with the caudal hematopoietic tissue, is another site of *tfec* expression; [37]) and die after approximately 12 days, apparently from lack of ability to feed. Potential postembryonic roles of *tfec* remain unclear, but data from mosaic adults and a single adult escaper suggest that *tfec* is required for iridophores throughout the lifetime of the animal (Fig. S1).

Our mutants presented with a complete absence of iridoblasts and mature iridophores but also, surprisingly, delayed differentiation of both NC-derived and RPE melanocytes. While differentiation of xanthophores is also delayed in *tfec* mutants, we found that development of all non-pigmented derivatives (neurons, glia and skeletal components) is unaffected. This striking phenotype restricted to pigment cell specification is unique amongst the characterised zebrafish pigment mutants. Decades ago, it was proposed that melanocytes share a common origin with iridophores and xanthophores from a pigment-restricted precursor [10], which we would term a chromatoblast (Cbl). More recently, analysis of *ltk* expression in *sox10* mutants indicated the presence of an *ltk*+ precursor of pigment cells, consistent with the chromatoblast [11]. The *tfec* loss of function phenotype thus provides further support for the existence of a Cbl, transiently localised within the dorsal trunk of embryos between 18 hpf and 24 hpf ([14]; Fig. 8).

**Figure 8.**
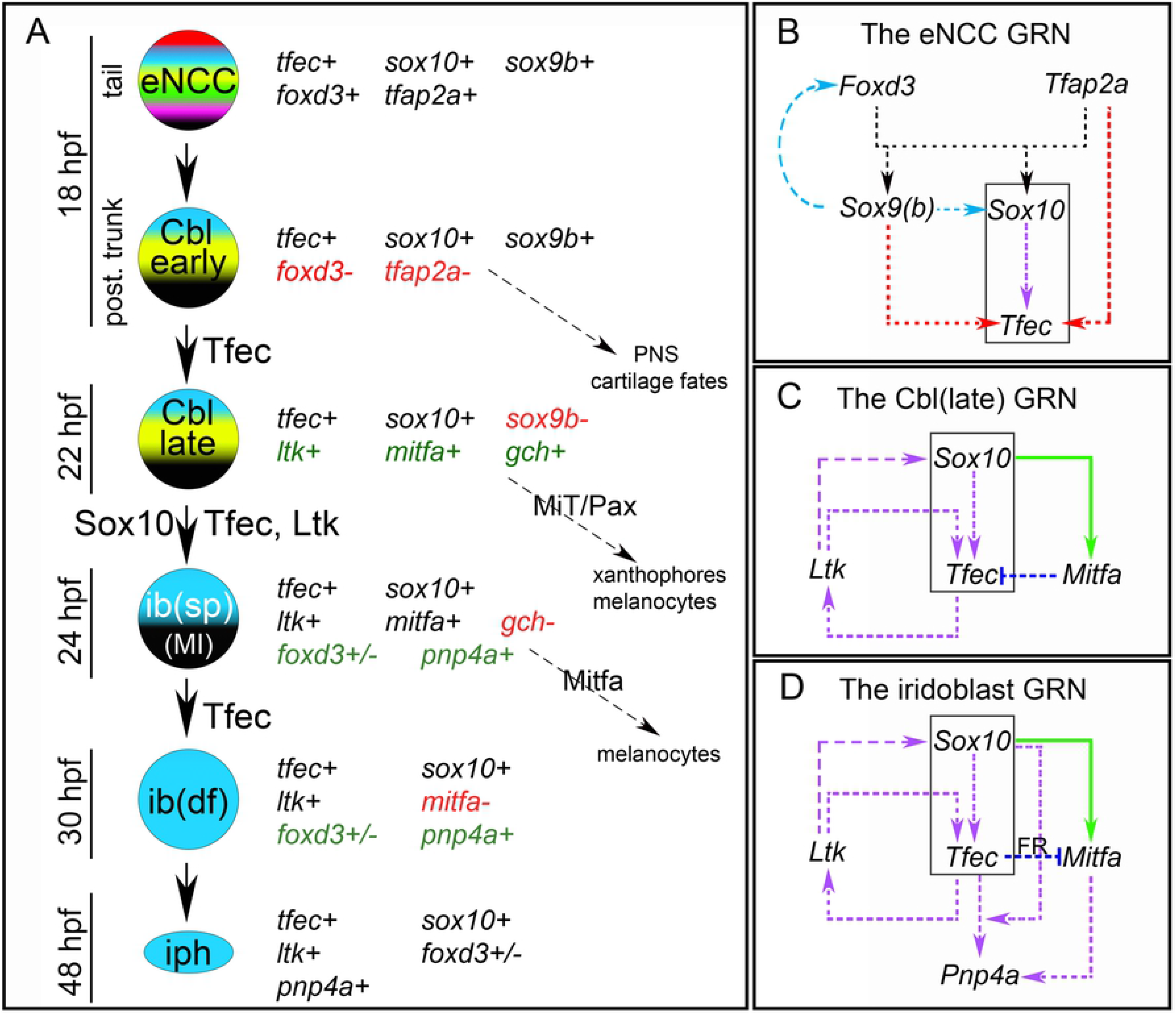
*tfec* is broadly expressed during progressive fate restriction of the iridophore lineage from eNCCs, and is required and sufficient to specify definitive iridoblast, ib(df), from the melano-iridoblast, ib(sp). (A) Schematic representation of partially restricted iridophore progenitors during development, along with the expression characteristics and potential fate choices of each ([14], this work). *tfec* is initially co-expressed with eNCC specification factors, which gradually become downregulated (red font), while lineage-specific factors become upregulated (green font). Proteins indicated on the arrows are considered important for the respective fate restriction step. (B) The position of *tfec* in the GRN guiding vertebrate NCC induction. Dashed arrows indicate interactions which could be either direct or indirect. In multipotent NCCs, Tfap2a and Foxd3 redundantly activate *tfec* expression (red arrows; this work), which is later maintained in the iridophore lineage by Sox10 (purple arrow; [14]). This activation occurs independently of previously described co-regulation of zebrafish *sox9b* and *sox10* expression by Tfap2a and Foxd3 (black arrows; [46]), and is unaffected by potentially conserved activation of *sox10* and *foxd3* by Sox9b (blue arrows, as shown in chick; [45], [47]). (C) Following transition of Cbl early to Cbl late, *tfec* expression is supported by positive feedback interactions between Sox10, Ltk and Tfec, as described in Petratou et al., 2018 [14]. Sox10 directly activates *mitfa* expression (solid green arrow; [23]), which during early Cbl specification is co-expressed with *tfec* [14], inhibiting its expression to bias progenitors towards the melanocyte fate. (D) As the lineage progresses past the Cbl, into the ib(sp), ib(df) and iph stages, factor R (FR) mediates Tfec-dependent downregulation of Mitfa (dark blue edge), and expression of the marker *pnp4a* becomes prominent [14]. In (B-D) the black box outlines the core connecting the three networks. MiT, microphthalmia family transcription factors; PNS, peripheral nervous system.

Based on our findings, *tfec* is the first zebrafish gene identified that specifically affects all chromatophore lineages without appearing to affect any non-pigment NC derivatives. While the key NC transcriptional regulator, *sox10,* is required for each of the three pigment cell lineages, loss of *sox10* function additionally results in strong reductions or absence of all peripheral glia cells and NC-derived peripheral neurons ([17], [20], [21], [23]). Furthermore, this requirement of pigment cells for *sox10* manifests itself apparently in different ways; *sox10* is required for *mitfa* expression in late Cbls, and *mitfa* can promote melanocyte fate in the absence of *sox10* if misexpressed [23]. In contrast, *sox10* mutants still strongly express *tfec* (as well as *ltk*) within trapped Cbls, yet fail to produce Ib(sp) [14]. We show here that in *tfec* mutants the ability to express the lineage markers of melanocytes and xanthophores is delayed but otherwise unaffected, whereas *ltk* expression is completely missing, even in Cbl(late). Thus, Tfec is necessary for both correct development of the Cbl(late) and for Ib(sp) specification from such a cell.

In order for *tfec* to be considered the iridophore master regulator, the gene must not only be necessary, but also sufficient for iridophore specification. Our data are strongly suggestive of *tfec* being required for iridophore development, as *mitfa* is for melanocytes (Fig. 8). Injection of wild-type *tfec* cDNA under its own promoter could rescue, albeit only partially, the iridophore phenotype in *tfec* mutants, with rescued iridophores in the eye observed more often than in the trunk. Whilst these results show that Tfec is sufficient to rescue iridophore specification in tfec mutants, fulfilling the conditions for a master regulator gene, gain of function experiments will be required to determine if the mechanism by which Tfec functions in iridophore specification is analogous to that of Mitfa, which can trigger melanocyte development in the absence of *sox10* by independently triggering a feedback loop and expression of melanogenic genes ([15], [23]). Further work to define direct and indirect transcriptional targets of Tfec, as well as any other transcriptional co-regulators, similar to the role of Tfap2a for Mitfa [25], is needed to understand the mechanistic details of iridophore fate choice. RNA-sequencing and ATAC-sequencing assays on iridophore precursors would be invaluable to determine, in an unbiased manner, appropriate candidates.

Another aim of the work presented here was to extend the GRN surrounding *tfec* beyond the iridophore lineage by examining its regulation in early NCCs ([14], [37]). Initiation of *tfec* expression in the early NC is not affected by loss of *tfap2a*, *foxd3* nor *sox10* alone. Notably, loss of *sox9b* alone caused an apparent delay in induction of *tfec* expression in posterior NC yet, despite this alteration in the pattern, expression remained strongly present in the progenitor population.

Sox10 and Sox9b have previously been reported to act redundantly during zebrafish development [15], [20]. Our data, however, demonstrates that *tfec* expression in double *sox10;sox9b* mutant embryos is still strongly activated in the NC. It remains to be shown whether loss of function of a single or of both *sox10* alleles modifies the degree of *tfec* expression delay noted in homozygous *sox9b* mutants.

Double mutants for *tfap2a;foxd3* have been shown to eliminate induction of NC [42]. Redundant activities of Tfap2a and Tfap2c are required for NC induction and development of other non-neural ectoderm derivatives in zebrafish embryos ([42], [43]). Consistent with this, we found that *foxd3* and *tfap2a* are redundantly required for induction of *tfec* expression in early premigratory NCCs (Fig. 8B). In this context, it is interesting that *tfec* was recently identified as a likely direct target of Tfap2a/2c, through analysis of gene expression changes in mutants combining different numbers of *tfap2a* and *tfap2c* mutant alleles [44]. The same study demonstrated the functional compensation of Tfap2a by Tfap2c, since in *tfap2a* mutants just a single copy of *tfap2c* was sufficient to maintain *tfec* expression at WT levels and rescue NC specification. Our data support the above finding, showing that transcriptional regulation of *tfec* via Tfap2 transcription factors is independent of them first activating *sox10* or *sox9b* ([45]–[47]) (Fig. 8B). Moreover, our assays indicate that disruption of Foxd3 activity alongside Tfap2a counteracts functional compensation by Tfap2c. Further work will be required to assess whether this is due to a role for Foxd3 in Tfap2c expression.

Our data formally establishes *tfec* as a member of the GRN governing maintenance of NCC progenitors [39]. Furthermore, our data provides the first evidence for Tfec function in those early progenitors, since we show it is required to drive early expression of *ltk* (in Cbl(late)), as well as fate specification of the iridophore lineage from these multipotent progenitors.

Although *tfap2a* and *foxd3* act redundantly to activate *tfec* expression in early NCCs, they also present with divergent ongoing effects in pigment lineages. Single mutations in *tfap2a* and *foxd3* affect the melanocyte and iridophore lineages, respectively ([26], [27], [48]), likely in a manner dependent upon distinct regulatory interactions with Mitfa. While *tfap2a* and *mitfa* work in parallel to promote melanocyte differentiation [25], *foxd3* has been suggested to repress *mitfa* transcription ([12], [49]), at least in some contexts [14]. In the absence of *foxd3*, iridophore numbers are reduced in a manner that is at least partially *mitfa*-dependent, and marker analyses and lineage-tracing experiments support the existence of a bipotent melanocyte-iridophore (MI) precursor, the fate of which is influenced by this *foxd3/mitfa* interaction [13]. Our findings that *mitfa* represses *tfec* expression in melanoblasts, and that *tfec* mutants have increased melanocytes, indicate that maintenance of *tfec* activity is also key to the melanocyte versus iridophore cell fate decision. Notably, Mitfa and Tfec have the potential to physically interact as a heterodimer ([50], [51]), which adds an additional layer of complexity when attempting to elucidate the mechanism of cell fate choice. Interestingly, despite the similarity between *tfec* and *ltk* mutant iridophore phenotypes, *ltk* mutants do not show an analogous late increase in melanocytes [11].

Specification and differentiation of the third pigment cell type of zebrafish, the xanthophore, has been shown to depend upon the paired-box transcription factors Pax3 and Pax7a/b ([52], [53]). Intriguingly, interactions between Pax3 and Mitf have been demonstrated in mammal melanocyte development [54]. At least some zebrafish xanthoblasts express *mitfa* [55] and we show that loss of *tfec* delays xanthoblast migration, raising the question of whether interplay between Mitfa and Tfec might be important for xanthoblast fate choice. Furthermore, it will be of interest to examine potential interactions between not only Tfec and Mitfa, but also between MiT and Pax transcription factors during chromatoblast diversification.

To conclude, our study contributes to deepening our understanding of the molecular basis of NC and pigment cell development in zebrafish, as well as the process of progressive fate restriction of multipotent stem cells. Detailed assessment of the diversity of pigment progenitor states in these embryonic stages is needed to test the hypothesis of a tripotent chromatoblast. Furthermore, focused effort on the (redundant) roles of transcription factors in xanthophore development will be decisive in understanding pigment cell fate choice in zebrafish.

## Materials and Methods

### Ethics statement

This study was performed with the approval of the University of Bath ethics committee and in full accordance with the Animals (Scientific Procedures) Act 1986, under Home Office Project Licenses 30/2937 and P87C67227, and in compliance with protocol AM10125 approved by the Institutional Animal Care and Use Committee of Virginia Commonwealth University.

### Zebrafish Husbandry

All embryos were obtained from natural crosses. Staging was performed according to Kimmel et al (1995) [56]. WIK or NHGRI-1 wild-type embryos served as controls as indicated in each figure. The following mutant lines were examined: *sox10^t3^* [18], *foxd3^zdf10^* [27], *sox9b^fh313^* ([57], [58]), *mitfa^w2^* [19], *ltk^ty82^* [11], *tfap2a^ts213^* [59]. Adult fish were maintained in accordance to official guidelines: water temperature: 28 °C, water pH: 7.4, conductivity: 800 mS, ammonia concentration: <0.1 mg/L, nitrite concentration: 0 mg/L, nitrate concentration: <5 mg/L, water hardness: 80-120 mg/L. The dark/light cycle was set at 10/14 hours. The adult stocking density was 5 fish/L, and the larvae stocking density <100 fish/L.

### CRISPR mutagenesis

Target sequences for CRISPR/Cas9 mutagenesis were identified using version 1 of the CHOPCHOP webtool (http://chopchop.cbu.uib.no/; [60]). The template for synthesis of the guide RNA (gRNA) was generated using a previously described PCR method [61]. Primer sequences are provided in Table 1. The *tfec^ba6^* and assorted *tfec^vc^* alleles were generated independently as follows:

**Table 1.**
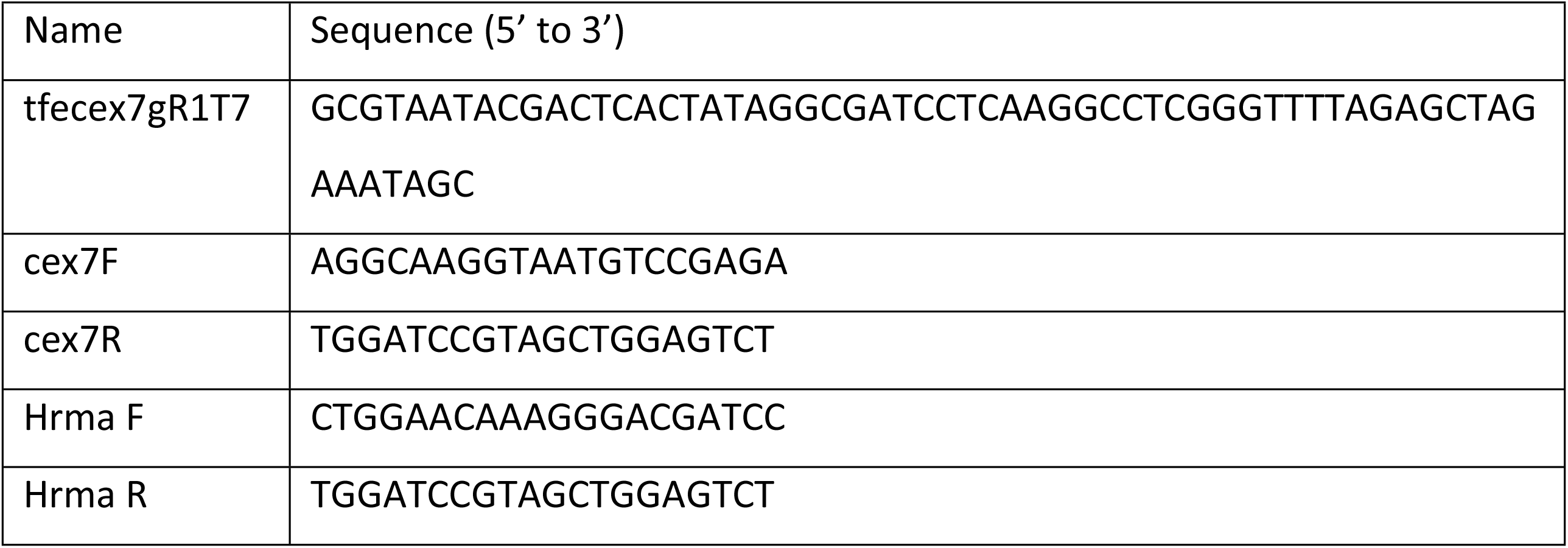
Oligonucleotides used in this study

### *tfec*^ba6^

280 pg of CRISPR guide RNA mixed with 700 pg *Cas9* mRNA per embryo were injected at the flat cell stage. RNAs were diluted in RNase-free water. Approximately 200 injected embryos were raised, from which 30-40 pairs were in-crossed to screen for germline transmission. Adults that transmitted mutant alleles were separated and outcrossed wild-type fish of the WIK line to generate F1 offspring. F1 siblings (which conceivably could carry different mutant alleles) were in-crossed to identify mutation carriers. 3 fish were identified, which were outcrossed again to WIK fish and the resulting F2 generation raised. For all experiments the F2 of one of the three identified F1 fish were used. Screening was based on iridophore phenotype and confirmed by a High Resolution Melt Analysis (HRMA) assay for molecular screening (see below). To characterise the mutations, F1 adult genomic DNA extracted by swabbing and genomic DNA extracted from F2 embryos were sent for sequencing.

### *tfec*^vc^ *alleles*

CRISPR guide RNA was synthesized using the MEGAshortscript T7 Transcription Kit (Invitrogen, Cat# AM1354) and purified using the miRvana miRNA Isolation Kit (Invitrogen, Cat# AM1560) as described by [62]. Capped *Cas9* mRNA was generated from the plasmid pT3TS-nCas9n [62], a gift from Wenbiao Chen (Addgene plasmid, Cat# 46757) using the mMESSAGE mMACHINE T3 Transcription Kit (Invitrogen, Cat# AM1348). *Cas9* mRNA and *tfec* exon 7 sgRNA were each diluted to 100 ng/μl for microinjection into one-cell embryos of the NHGRI-1 strain [63]. A fraction of the injected set was sacrificed for genomic DNA preparation [64] to evaluate the efficacy of the guide RNA using the primers cex7F and cex7R, followed by restriction digest with *StuI,* which cuts in the target sequence. The remaining injected embryos were raised to adulthood and intercrossed or mated to wild-type (NHGRI-1) partners. F1 carriers were identified using the PCR digest assay above, and undigested PCR products (representing mutant alleles) were purified and sequenced.

## High resolution melt analysis

High Resolution Melting (HRM) Software V3.0.1 (Thermo Fisher Scientific) was used to detect and amplify differences between the melting temperature of 150-200 bp q-RT PCR amplicons generated from reference wild-type (WT) samples versus mutagenized embryos or adults. To perform q-RT PCR for HRMA, template genomic DNA was extracted using the KAPA Express Extract Kit (Sigma-Aldrich; Cat# KK7103) according to manufacturer’s instructions, and was diluted to 8 ng/μl. Amplification reactions were set up as per manufacturer’s instructions using KAPA HRM FAST reagents (Sigma-Aldrich; Cat# KK4201) and primers designed according to the relevant recommendations (Table 1). Following amplification, a continuous melt curve was generated by increasing the temperature from 60 °C (1 min) to 95 °C (15 sec) in 0.3 °C /sec increments. To detect CRISPR/Cas9-mediated mutagenesis, at least 8 WT reference samples were included in the analysis.

## Chromatophore counts

Melanocyte counts were performed at 30 hpf and at 4 dpf on anaesthetized or fixed embryos under transmitted light. Embryos at 4 dpf were treated with 2 μM melatonin directly prior to counting. To count iridophores, PTU-treated embryos were observed under incident light. Pigment cell counts were made under a Zeiss Axio Zoom.V16 fluorescence stereo zoom microscope.

## Cloning and rescue by plasmid microinjection

The full-length coding sequence of *tfec* was subcloned in-frame into the Gateway 3’ Entry vector p3E-2A-FLAG-pA to make p3E-2A-FLAG-tfec-pA. A multisite Gateway LR+ cloning reaction was then carried out with this plasmid along with Tol2 Kit destination vector pDestTol2pA2 and entry vector pME-mCherry-no stop [65] and entry vector p5E-tfec2.4 [40]. Following bacterial transformation, correct clones were identified by restriction digest of miniprep cultures.

The resulting plasmid (designated pDestTol2pA2-tfec2.4/mCherry-nostop/2A-FLAG-tfec-pA) at 25 ng/μl was co-injected with Tol2 transposase mRNA at 25 ng/μl into embryos from an intercross of heterozygous *tfec^vc60^* carriers. Homozygous mutant embryos were identified between 48 and 72 hpf and scored at 96 hpf on a stereo dissection microscope under incident light for the presence of iridophores.

## Transcript detection in whole-mount embryos

Detailed information on the preparation of generic materials and the protocols for performing chromogenic WISH as well as multiplex fluorescent RNAscope can be found in Petratou et al. (2017) [66]. Probes used for chromogenic WISH were *sox10* [18], *foxd3* [67], *ltk* [11], *pnp4a* [13], *aox5* [55], *gch2* [55], *pou3f1/oct6* [68], *dlx2a* [69], *mitfa* [19] and *tfec* (NM_001030105.2, [14]). For multiplex RNAscope, the following probes were used: *ltk* (ACD, Cat# 444641), *tfec* (ACD, Cat# 444701).

Embryos were observed and imaged, and the Pearson’s chi-squared test was used, as previously described [14], to process and statistically analyse results, to test the hypothesis that altered phenotypes correlated with homozygosed mutations. *tfap2a* and *foxd3* mutant embryos were identified using previously described genotyping protocols ([27], [70]) following imaging and preparation of genomic DNA [64].

## Alcian blue staining and immunohistochemistry

Larvae from an intercross of *tfec^vc60^* heterozygous adults were sorted at 4 days postfertilization based on the iridophore phenotype, and then were fixed in separate tubes overnight in 4% PFA. Alcian blue staining was then carried out essentially as described [71]. Samples were imaged using an Olympus SZX12 stereomicroscope with DP70 camera.

Immunohistochemistry was carried out as previously described [68]. Primary monoclonal antibodies against HuC/D (Molecular Probes) and Pax7 (Developmental Studies Hybridoma Bank) were used at 1:500 and 1:20 respectively, and goat anti-mouse secondary antibodies conjugated to Alexa 568 or Alexa 488 (Molecular Probes) were each used at 1:750 dilution. For combined *tfec* WISH/Pax7 IHC, the Pax7 antibodies were added simultaneously with the anti-Fab fragments [72]. Samples were imaged on a Zeiss Axio Imager.M2 compound microscope with Axiocam 503 colour camera, processed using ZEN software and Adobe Photoshop CC 2018 and 2019.

## Supplementary file legends

**Figure S1. Loss of *tfec* affects adult iridophore pigmentation.** Compared to WT adult (A), G0 *tfec* crispant (mosaic) adult shows patches of iridophore loss on eye and flank (B). When two *tfec* CRISPR G0 founder fish were mated, almost all of the offspring lacking iridophores died as larvae, but one escaper survived to adulthood and the absence of iridophores persisted (C). Of the homozygous *tfec^vc60^* embryos injected with a Tol2 transposon containing the *tfec* promoter and cDNA and Tol2 transposase mRNA, one survived to adulthood and displayed partial rescue of adult iridophore pigmentation (D). Scale bar: A-D: 0.5 cm; D: 0.35 cm.

**Figure S2. At 24 hpf, *tfec* expression is found in trapped NC precursors in *tfap2a* homozygous mutants.** (A) Wild-type *tfec* expression is detectable in dorsally positioned, and ventrally migrating Ib(sp) along the anterior trunk (asterisk). *tfec*+ eNCCs are located in the posterior of the embryo (region between the arrowheads). (B) *tfec*+ eNCCs are detectable along the anterior and posterior trunk and tail of a *tfap2a* homozygous mutant sibling (area marked by black arrowheads). Expression is also strongly detected in cranial NCC derivatives (red arrows), which in the WT have turned down *tfec* expression. Insets show a higher magnification view of the WT (A) and mutant (B) posterior trunks. Lateral views, head towards the left. Scale bars: 50 μm.

**Table S1: Additional information on the assessment of live embryonic phenotypes.** For melanocyte counts at 4 dpf, to derive the average along the lateral stripe, the cells along both stripes of each embryo were independently scored. Presented p-values derived from unpaired two-tailed t-test between WT (or heterozygous for the melanocyte counts) and the genotype corresponding to each row. DS, dorsal stripe; DT, dorsal trunk; H, head; LS, lateral stripe; MP, migration paths; VS, ventral stripe; VT, ventral trunk.

**Table S2: Statistics of loss of function experiments.** The Pearson’s chi-squared test for goodness of fit is used to calculate the likelihood of a non-WT phenotype, which is consistently present in a number of embryos (1^st^ sub-column of each of the 4 developmental stages) within a batch of WT, heterozygous and homozygous mutant siblings, correlating with homozygosity of the mutant allele in those individuals. Due to the recessive nature of investigated alleles, 25% of embryos in each batch are expected to be homozygous mutants. Therefore, the p-value derived from the chi-squared test indicates whether the number of individuals with an alternative phenotype conform to the expected 25% (null hypothesis), with any deviation being attributable to random chance (p > 0.1), or whether the numbers deviate significantly from the expected (p ≤ 0.1; null hypothesis is rejected). Where observed phenotypes did not significantly correlate with expected Mendelian ratios, i.e. where there is unlikely to be a mutant phenotype, the corresponding counts and p-values are red and bold. For the data on *sox10^t3/t3^*; *sox9b^fh313/fh313^* double mutants, the orange cells indicate how many embryos in the sample showed the known *sox9b^fh313/fh313^* phenotype, the green cells indicate number of embryos with the *sox10^t3/t3^* phenotype and blue cells indicate unexpected alternative phenotypes, likely owed to double loss of function.

